# Ancestral genetic components are consistently associated with the complex trait landscape in European biobanks

**DOI:** 10.1101/2023.10.04.560881

**Authors:** Vasili Pankratov, Massimo Mezzavilla, Serena Aneli, Daniela Fusco, James F Wilson, Mait Metspalu, Paolo Provero, Luca Pagani, Davide Marnetto

## Abstract

The genetic structure in Europe was mostly shaped by admixture between the Western Hunter-Gatherer, Anatolian Neolithic and Steppe’s Yamnaya ancestral components. Such structure is regarded as a confounder in GWAS and follow-up studies, and gold-standard methods exist to correct for it. However, it is still poorly understood to which extent these ancestral components contribute to complex trait variation in present-day Europe.

In this work we harness the UK Biobank to address this question. By extensive demographic simulations and incorporating previous results obtained using the Estonian Biobank, we carefully evaluate the significance and scope of our findings.

Heart rate, platelet count, monocyte percentage and many other traits show stratification similar to height and pigmentation traits, likely targets of selection and divergence across ancestral groups. The consistency of our results across biobanks indicates that these ancestry-specific genetic predispositions act as a source of variability and as potential confounders in Europe as a whole.

## Introduction

In order to uncover the genetic basis of complex traits, a large amount of data to be analyzed in extensive Genome Wide Association Studies (GWAS)^1,2^ has been collected in nation-wide population-based Biobanks^2–7^. Despite including predominantly individuals of European ancestry, which poses well-recognized limitations^2,8–10^, these cohorts still contain many stratifications not connected with the trait of interest, such as socio-economic disparities, geographic factors and, most importantly, inherent genetic structure. A plethora of methods^11–14^ have been developed to correct for these unwanted sources of variance that might bias GWAS discovery. Indeed even the finer cases of population structure present in national Biobanks ^15–17^ have been demonstrated to affect GWAS^18–23^ and, if not carefully addressed, hamper analyses following up on these results, such as Polygenic Risk Scoring^19,21,22,24–26^ and polygenic selection testing^27,28^.

Millennia of demographic expansions, migrations, and localized genetic isolation have indeed shaped a far from homogeneous genetic makeup for contemporary Europeans. Besides recent demographic history, predominantly impacting on rare variation structure, a series of admixture events from 8000 BCE to 1000 BCE have been foundational to the European genetic landscape. During that period, Anatolian Neolithic Farmers (Anatolia_N) and Steppe Yamnaya pastoralists (Yamnaya) joined local Western Hunter-Gatherers (WHG)^29–31^ with different times and modes across Europe, contributing with genetic components that had hitherto evolved separately for up to 20,000 years^32^. Divergent phenotypes in these source populations have been previously described for a few traits using polygenic scoring of ancient samples^33–35^, or looking at specific trait-informative Single Nucleotide Polymorphisms (SNP)^36–38^.

These ancestral components can explain a large part of the genetic gradients across Europe^30^ and, as a result, are indirectly adjusted for in gold-standard GWAS procedures. In this work, we aim to identify which complex trait variations across the largest European biobank can be explained by the stratification of these ancestral components and consequently to characterize the phenotypic predispositions conferred by these components. We previously attempted to quantify their differential contribution to the contemporary landscape of complex traits in 35,000 individuals from the Estonian Biobank (EstBB)^39^. Our findings provided a first picture to describe the association of ancestral populations to present day traits; however, it remained unclear whether our conclusions could be applied to the broader European population. In addition, we could not decisively address the differences in the resulting signal when running the analysis over local candidate regions or the whole genome. Finally, the study lacked a meticulous simulation testing the limits of the adopted statistic in answering our question.

Here we overcome these limitations by analyzing a total of 53 complex traits in 50,000 UK Biobank^3^ (UKBB) donors taken as representative of the Western European metapopulation. We validate the ancestry-trait associations found as European-wide signals, find new ones, and explore the strengths and limitations of our approach with thorough simulations using SLiM^40^.

## Results

### Framework overview

Our approach is based on *covA*, a measure of relative genetic similarity of an individual from a contemporary population to the distinct ancestries (e.g. WHG; Anatolia_N; Yamnaya) that contribute to the genetic makeup of that population. This metric was introduced in our previous work^39^ for the analysis of complex traits in EstBB and is ultimately a covariance between allele dosage in a contemporary individual and a given ancestral population, with respect to the contemporary and ancient average frequencies. We regress each complex trait *t* in the present-day dataset on the *covA* for each ancestry *p* so that for each individual *i*:

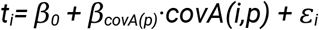

where the slope *β*_*covA(p)*_ quantifies the association between each ancestry *p* with the trait *t*.

In order to reduce the influence of genomic confounding factors, we compute *CovA* restricting to variants belonging to 20kb genomic regions around SNPs that have been previously associated with the trait of interest through GWAS, i.e. Trait-Associated Genomic Regions (TAGR, see Methods). Note that besides allowing us to identify TAGRs we avoid the use of GWAS-derived summary statistics. Using these summaries as weights in follow-up analyses, especially when summed across the whole genome, has been shown to produce results difficult to interpret in populations that are even subtly genetically differentiated from the ones where the GWAS was run^22,24,28,41^.

### Simulations

To explore *covA* strengths and limits, we performed 2370 demographic simulation runs in SLiM^40^ mimicking the major demographic events in the British population history and applying different selection regimes (various strengths of stabilizing selection, differences in trait optimum between ancestral populations, post-admixture selection). Each simulation run resulted in genotypes at 1000 unlinked loci of 20kb (portraying TAGRs) together with fictitious trait values for around 7K unrelated present-day individuals and corresponding simulated WHG, Anatolia_N and Yamnaya samples. See Methods and Supplementary Methods sections for further details.

To assess the performance of *covA*, we first ranked the three reference populations (equivalent to the sampled ancient genomes used to calculate *covA*) for their mean genetic value (GV), that is the expected phenotypic value without considering the environmental deviation. Therefore in each run *p=1,2,3* is the population with the highest, the median, and the lowest mean GV respectively. In scenarios of directional selection *p=1* corresponds to the ancestry experiencing a positive shift in trait optimum. We then tracked the slope obtained when regressing the simulated trait on *covA(1)* i.e. *β*_covA(1)_.

We started by verifying whether *β*_covA_ depends on the difference in the trait’s mean genetic value between reference populations. On average, higher *β*_covA(1)_ values are observed as the difference in GV between ancestries *p=1* and *p=2* increases (**Figure 1A**). In other words, *covA* is the more predictive of the trait value in present-day individuals the more genetically differentiated are the reference populations at TAGRs. On the other hand, we see no independent effect of heritability, polygenicity or strength of stabilizing selection on the relationship between *β*_covA_ and genetic value differentiation (**Figure S1**).

**Figure 1.**
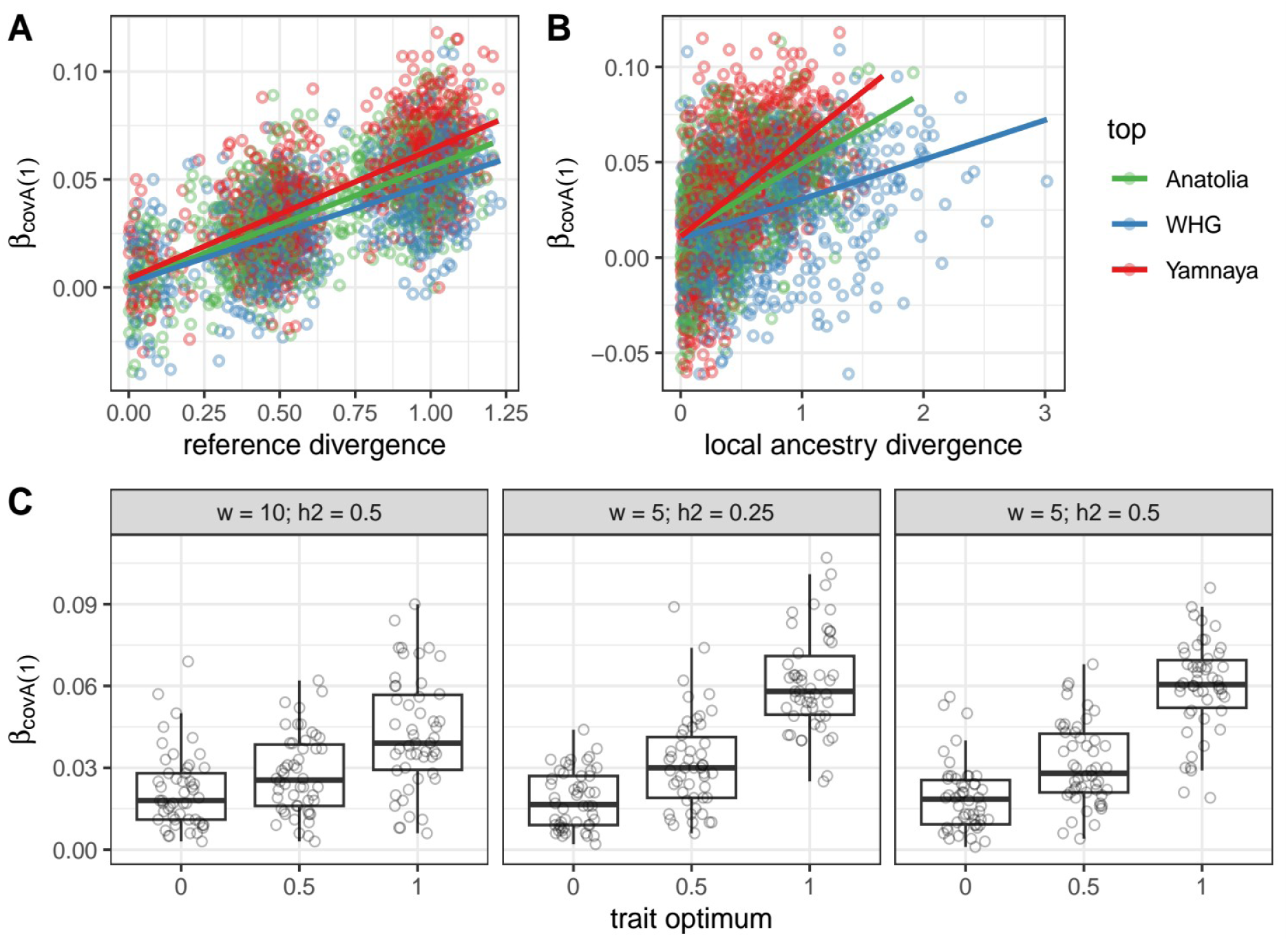
Simulation Results. (A) Effect on β_covA(1)_ of reference populations differentiation, defined as the difference between the highest and the second highest mean genetic values among the three ancestries, normalized by trait standard deviation in the present-day population. The trait is measured in arbitrary units, however, by design its standard deviation is mostly within the range between 0.9 and 1 (**Figure S6**). (B) The same as (A) but plotting on x-axis local ancestry’s mean genetic values instead. (C) β_covA(1)_ values as a function of trait optimum in the ancestry with the highest genetic value (p=1). The x-axis indicates the trait optimum in this ancestry, while other ancestries have an optimum set to 0. Three sub-panels present results of simulations with different fitness function SD (ω) and heritability (h^2^) values. Each boxplot is based on 50 individual simulations. Scenarios with more than 50 simulation runs were randomly downsampled. The boxes show 25^th^, 50^th^ and 75^th^ quantiles, while the whiskers show values within 1.5 times the interquartile range (IQR).

Notably, we can reliably identify the reference ancestry *p=1* as the one with the maximal *β*_covA_ as long as it is sufficiently differentiated from the ancestry *p=2*: the true-positive rate is 78-87% if the difference between GV is within 0.25-0.5 of the trait’s SD in the present-day population (**Figure S2A**). Note, however, that *β*_covA_ values are negatively correlated between ancestries (**Figure S3**), meaning that high absolute *β*_covA_ for one ancestry will be often complemented by high *β*_covA_ for at least one other ancestry but with an opposite sign.

We next asked if *β*_covA_ can be informative about differences in trait optima between ancestries. We hypothesize that drift under stabilizing selection with the same optimum in all ancestries would result in weaker GV differentiation and lower *β*_covA(1)_ as compared to scenarios with optimum shift in one of the ancestries. Although we do observe such a trend (**Figure 1C, S4**), rather high values of *β*_covA(1)_ (97.5 percentile of 330 simulations being equal to 0.048, see **Table S1**) can also be obtained in simulations with constant trait optimum under various levels of heritability and stabilizing selection.

Finally, we checked whether *β*_covA_ can be informative about differential genetic contribution of ancestral populations to present-day trait variation. Intuitively, this can be measured as the expected GV of a present-day individual with all trait-affecting loci inherited from a specific ancestry. This value might deviate from the reference population’s average GV due to post-admixture drift and selection. As can be seen in **Figure 1B** and **Figure S5**, *β*_covA_ remains informative about the difference in local ancestry GV between ancestries p=1 and p=2 but the true-positive rate of identifying the local ancestry with the highest GV is lower (57-78% if GV difference is within 0.25-0.5 of the trait’s SD in the present-day population; **Figure S2B**). The lowest sensitivity is observed for the WHG local ancestry likely because of its low contribution to the present-day population (12%) and hence stronger post-admixture drift. However, as exemplified by scenarios **E** and **F** in **Figure S5**, when the reference populations had the same trait optimum, our approach is blind to differences in local ancestry genetic value arising from post-admixture drift (**E**) or selection (**F**). Full simulation results are reported in **Table S2**.

### Association with UK Biobank complex trait landscape

We applied the same approach to 50,000 unrelated individuals of European descent from the UKBB^3^, so as to match the order of magnitude of our previous EstBB analysis^39^. The reference groups representing WHG, Anatolia_N and Yamnaya (here defined as genetic components rather than cultures) are identified starting from a core hand-picked set of ancient samples from the Allen Ancient DNA Resource^42^, then expanded with samples with similar date and genetic principal components. With this process, we selected 95 WHG-, 118 Anatolia_N-, 83 Yamnaya-like ancient genomes, see list in **Table S3, Figure S7**.

We identified TAGRs for each trait exploiting GWAS Catalog^43^ hits for congruous traits (see methods for details). Indeed, TAGRs were enriched in SNP heritability for the corresponding trait in almost all cases (stratified LD score regression^44^, nominal P-value<0.05, see **Figure S8A**). We then computed *covA*s for each ancestry on these regions and regressed phenotypic values for each trait on them. Notably, to capture any possible physiology-related, geographic and socio-economic confounder that might produce spurious false positives, we also include in this regression a large set of covariates: sex, age, age2, sex×age, sex×age2, genome-wide *covA* for two ancestries (capturing all *covA* variability), latitude, longitude, UKBB assessment centre, Townsend deprivation index (TDI), qualifications, age when completed education. In addition, for binary or ordinal traits we fitted logistic and ordinal logistic rather than linear models, and assessed the estimated Odds Ratio (OR).

An independent model was built for each of the ancestries as our simulations showed that a genetic value shift in a single ancestry could explain significant *β*_*covA*_ in all ancestries (**Figure S3**), and that the ancestry with the maximal absolute coefficient (*max(*|*β*_*covA*_|)) is likely to be the most divergent one. As seen in **Figure 2A**, 32 out of 53 traits tested show at least one significant *β*_*covA*_ (or OR_*covA*_) at 5% Benjamini-Hochberg False Discovery Rate (FDR). Affected traits include several biological domains and will be addressed in the next section. Notably, no trait exhibits a *max(*|*β*_*covA*_|) > 0.048, which based on our simulation results would have been a strong indication of pre-admixture selection.

**Figure 2.**
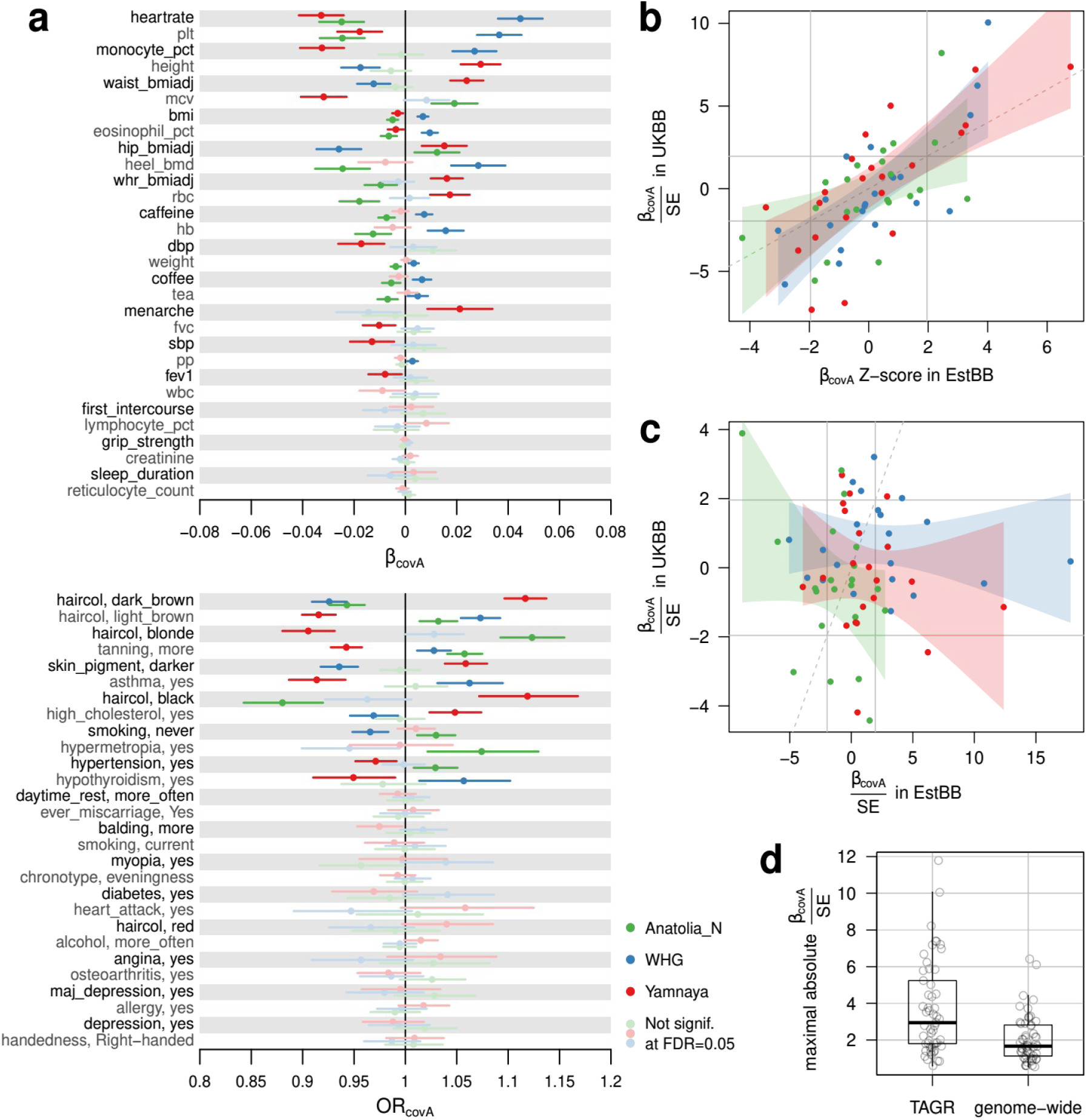
Trait-ancestry associations in UKBB and across Biobanks. **(A)** CovA effect size (β_covA_ or OR_covA_) estimates together with their 95% confidence interval, for each ancestry and for 53 traits analyzed. For each trait, covAs are computed on TAGRs and used in independent models as trait regressors together with several covariates. covAs not adding significant information according to double-sided coefficient p-value at 5% FDR are shown in pastel colors. **(B)** CovA test statistics compared between UK Biobank and Estonian Biobank, with CovA computed on TAGRs. Z-scores in X axis are extracted from the original paper^39^, and computed differently than β/SE. Shaded areas represent linear regression slope confidence intervals for each ancestry independently, the solid gray lines mark nominal significance boundaries for β_covA_ while the y=x line is represented as dashed. **(C)** Same as **(B)** but here CovA is computed genome-wide. **(D)** Maximal absolute CovA test statistics in the TAGR and genome-wide analyses. Each dot is a trait.

For 21 traits already explored in the Estonian Biobank (EstBB)^39^ we compared the regression statistics obtained across biobanks, finding a remarkable consistency, see **Figure 2B**. Even though we refrain from assigning a p-value due to the plausible correlation across several trait pairs, we identify a Pearson’s correlation coefficient between biobanks of ρ=0.555, ρ=0.783 and ρ=0.750 respectively for Anatolia_N, WHG, and Yamnaya ancestries.

We repeated the analysis by evaluating the coefficients of *covA*s computed on the whole genome, as opposed to only TAGRs. These were previously used as genomic control covariates, thus allowing us to only keep physiology-related, geographic and socio-economic factors as covariates this time. In this setting we can still see 15 out of 53 significant signals, see **Figure S9**, seemingly suggesting that the global similarity with a specific ancestry could be associated with a certain trait. However, the signals are often discordant and weaker on average: again we refrain from conducting a formal test due to correlation between traits but the trend is appreciable in **Figure 2D**. Moreover, the consistency across biobanks is completely absent if we compare genome-wide *covA*s, see **Figure 2C**, suggesting that these signals are likely resulting from yet unaccounted confounders rather than deriving from the trait inherent biology. Indeed, although likely harboring variants contributing to the polygenic traits analyzed, the remaining non-TAGR genome was often depleted in trait heritability (see **Figure S8B**) but equally subject to confounders impacting at genomic level.

### Ancestry-specific diverging traits

We interpret significant *β*_*covA*_ as ancestry divergence in genetic value and designate the genetic component showing max(|*β*_*covA*_|) as the most divergent one. Note however, that inopposition to our simulations, where only one ancestry was divergent by design, the ancestry with max(|*β*_*covA*_|) might not be the only one that experienced a shift in GV.

The largest signals found in our analysis are divergences in resting heart rate, platelet count and monocyte percentages in blood which increase together with WHG ancestry similarity, see **Figure 2A**. As shown by our simulations, adaptation might not be necessary to observe such divergence. Nevertheless, given health-relevant changes in platelet count and heart rate variability during winter^45,46^, coupled with the complex heart rate reactions to cold exposure^47,48^, it is tempting to speculate about an adaptation to colder climates that WHG suffered when colonizing Europe soon after the last glacial maximum (after 17kya ^49^). A similar pattern of divergence can be observed for BMI, hemoglobin concentration, eosinophil blood concentrations and heel bone mineral density. Interestingly, some of these traits such as platelet count and BMI were found to be selected for lower levels during the early Neolithic^50^ thus supporting the idea of higher values being WHG-specific and possibly maladaptive in later periods, in favor of more recent incoming genetic components.

We replicate known signals of genetic divergence across ancestries for anthropometric and pigmentation traits^39^, most notably an association of Yamnaya ancestry with tall stature^33^ and large waist, but also with darker hair and skin pigmentation, surprisingly coupled with poor ease of tanning. These results corroborate the hypothesis that pigmentation decrease in Europe is rather due to post-admixture selection rather than impact of incoming Yamnaya migrations during the Bronze Age^34,50^.

Some red blood cell properties are also divergent, with lower corpuscular volume but higher cell count associated with Yamnaya ancestry, while the opposite is true for Anatolia_N ancestry. These properties are important readouts for B12, folic acid and moreover iron intake, potentially indicating adaptations to pastoralist or farming diets. In this and other cases consistent with findings in EstBB^39^ such as blood cholesterol levels, caffeine intake, but also previously mentioned traits (e.g. monocyte percentage, pigmentation), two ancestries are both highly significant for opposite trait values. This is expected due to the negative correlation between *β*_*covA*_ (**Figure S3**) but can also result from two ancestries diverging in opposite directions while the third ancestry having an intermediate genetic value. One last notable example of such a pattern is the frequency of asthma diagnoses, which appear to be positively associated with WHG ancestry and negatively with Yamnaya. Interestingly, the Yamnaya genetic component is also weakly associated with lower indicators of pulmonary function, such as FVC and FEV1.

In addition, we find weaker but significant associations of Anatolia_N ancestry with hypermetropia, already suggestive in EstBB^39^, and Yamnaya ancestry with lower blood pressure (both systolic and diastolic) and consequent reduced exposition to hypertension, which was not evident in EstBB^39^.

Other weakly divergent traits appreciable in **Figure 2A** include smoking status, age at menarche and hypothyroidism.

## Discussion

We applied *covA* to the largest European biobank to identify which complex trait population gradients can be explained by genetic similarity with one of the three main European ancestries: WHG, Anatolia_N, and Yamnaya. In our work, we found that relative similarity with a specific ancestral component significantly explains 32 out of 53 complex trait distributions. Our findings are globally consistent with an independent analysis conducted on the Estonian Biobank for 21 overlapping traits, thus confirming that these results should be taken as indicative of continental patterns rather than just regional ones.

This agreement between biobanks is lost when not restricting the analysis to TAGRs: whole-genome results were inconsistent both across biobanks and when compared with TAGRs results within the same dataset (**Figure 2, S5**). In other words: although we took measures to control for confounders by including multiple geographic and socio-economic variables into our models, only by enriching for biologically-relevant regions, and by controlling for genome-wide covA stratification, we could expose robust and consistent signals. This finding mirrors the known difficulty in discerning functional gene-trait associations from spurious correlations mediated by genetic structure ^19–21,25,27,28^, which are indeed independent between different populations and different biobanks with different recruitment strategies. In addition, while combining GWAS-derived effect sizes at genome scale is sensitive to subtle correction-surviving biases^26^, our approach does not rely on such methodology.

While we are confident about the robustness and the continental scope of the results presented in **Figure 2A**, the biological and evolutionary dynamics that can explain these associations remain rather complex to establish. We thus complemented the empirical analysis with extensive simulations to provide insights useful in interpreting our results.

Specifically, the simulations show that significant *β*_covA_ for at least one of the ancestries can be most directly interpreted as differences in genetic values between sampled reference populations. Although in our simulations such differences can be reached under scenarios with all reference populations having the same optimum, scenarios with optimum shift in one of the ancestries in general result in higher absolute *β*_covA_ (**Figure 1C**) for that ancestry. As the highest *β*_covA_ we observe on the real data are close to the 97.5th percentile of the *β*_covA_ distribution under equal optima in simulations, it is likely that the top of our list contains at least some traits that underwent directional selection in different ancestral populations. Indeed, our results highlight that resting heart rate, blood platelet count and monocyte percentage, bone mineral density and asthma diagnoses present divergence patterns compatible with adaptive selection on par with deeply scrutinized traits such as pigmentation and height, and deserve similar investigation.

Our simulation results also inform us about the potential limitations of our study. These mostly relate to interpreting *β*_covA_ as an indication of the differential genetic contribution of ancestries to trait variation in the present-day population. On the one hand, such contribution can be overlooked if genetic differentiation between ancestries post-dates the available aDNA samples used as references (**Figure S5E**). On the other hand, as most common alleles are expected to pre-date the split between the three ancestries, present-day individuals with higher GV for a given trait might show higher *covA* with an ancestry enriched for trait-increasing alleles, even without any direct genetic contribution from that ancestry. Thus, choosing and exhaustively including all relevant ancestral groups is crucial for a sound biological interpretation of *covA* signals. This limits the applicability of our approach to non-European populations for which the demographic history is less well understood and proper reference aDNA samples are missing.

In conclusion, the reported ancestry-trait associations are strongly indicative of ancestry-specific genetic predispositions, possibly due to pre-admixture selection, and under certain conditions indicate actual genetic contribution to the contemporary trait landscape. Especially for the most significant associations described above, researchers should assume the European population as fundamentally stratified due to its demographic history. This bias will need to be addressed in analyses following up on GWAS effect sizes, in order to avoid potential spurious results. This stratification acts therefore both as a source for variability in such traits and as a potential significant confounder for GWAS study across European cohorts.

## Methods

### Simulations

To explore the behavior of *covA* under different heritability, polygenicity, stabilizing selection and differences in trait optima between ancestral populations we performed hybrid simulations using a combination of SLiM^40^ and msprime^51^. Msprime was used to generate genetic diversity at mutation-drift equilibrium while SLiM was subsequently used to simulate the complex trait under a demographic model relevant for the British population (**Figure S10**) and various combinations of parameters specifying the trait’s genetic background and evolution. All in all, we ran 2370 simulations representing 135 simulation scenarios.

In step 1 run in msprime we simulated 1000 unlinked genomic regions each being 20 kb long with a uniform recombination rate of 1e-8 and uniform mutation rate of 1.25e-8 in a panmictic population with a constant effective population size of 14,000.

At the beginning of step 2, we picked one SNP per each region to have a causal effect on the complex trait and randomly assigned effect sizes to them which remained constant throughout the remaining part of the simulation. Genotype at these causal SNPs affects individual fitness (see below), thus allowing for directional and stabilizing selection.

The genetic value of an individual is calculated as a sum or derived allele dosage at those causal SNPs weighted by corresponding effect sizes. The trait value of an individual is calculated as the genetic value plus environmental noise value, sampled from a normal zero-centered distribution. Individual trait values are mapped to fitness using a normaldistribution with mean equal to population’s optimum and SD (designated as *ω*’ here) describing the intensity of stabilizing selection. The way we model individual fitness is inspired by the study by Yair and Coop, 2022^52^.

Each simulation run resulted in genotypes and trait values for 10K contemporary individuals (around 7K after filtering for relatives) and 100 individuals from each of the three reference populations (WHG, Anatolia_N, Yamnaya; no relatives filtering applied). Note that the reference populations were sampled at the time of the corresponding admixture event but were simulated as sister groups for the actual admixing populations with divergence time of 50 generations and the same trait optimum. The causal SNPs were ascertained based on the fraction of heritability of the trait they explain to mimic SNP discovery in GWAS, resulting in between 351 and 565 TAGR used to calculate covA (**Table S2**). See Supplementary Methods for further information.

### Sample selection and definition of ancestral groups

UKBB^3^ donors were selected among those identified as “caucasian” according Data-Field 22006 (which includes self-identification and genetic grouping), dropping those indicated in Data-Field 22018 (“genetic relatedness exclusions”), relatives with kinship coefficient >=0.04419 (third or lower degree relatives), and samples with more than one missing trait among those analyzed. We then extracted a subset of 50,000 samples equally divided between females and males for the following analyses.

Ancient samples were extracted from the Allen Ancient DNA Resource^42^ v52.2 (AADR) following the approach described in Marnetto et al. 2022^39^. After starting from a manually curated core set for each ancestral group, we expanded these core sets to other AADR samples according to distance in a 4-dimensional space defined by dating and first 3 Principal Components (PC). PCs were determined on a set of modern Eurasian and North African individuals west of Iran (included), where the ancient samples were projected; distance cutoff was defined by multi-dimensional ellipses with diameters equal to 3 core set SDs. See **Table S3** for a full list of ancestral group classifications, coordinates, dates and PCs.

We used phased, imputed genotype data for the UKBB set and intersected it with the ancient set, remaining with 1,087,822 genotyped variants in the merged dataset.

### Traits and candidate regions

Traits were selected to span across different domains and favoring high effective sample sizes, see **Table S4** for a complete list accompanied with the original UKBB Field ID. When multiple data points existed for an individual, the earliest one was considered. For each trait considered as continuous, individuals with values more distant than 4 IQRs from the upper or lower quartile were considered as outliers and discarded, then traits were standardized. Some traits were computed (waist/hip ratio, pulse pressure, caffeine,…), adjusted(waist and hip circumference, blood pressure,…) or log-transformed (BMI, creatinine,…), see the same table for such information.

Trait-associated genomic regions (TAGRs) were defined starting from GWAS Catalog^43^ data, downloaded on 09/05/2023. This resource collects small-scale variant hits which, depending on the original study, either are genome-wide significant (p-value < 0.5⋅10^-8^) or genome-wide suggestive (p-value < 10^-5^). We selected hits by matching a pattern to the trait definition given by the original study or by selecting a mapped Experimental Factor Ontology term, according to the **Table S5**, then defined 20kb windows centered on the selected hits and merged them, obtaining a set of TAGRs for each trait.

### Partitioned heritability

In order to assess TAGRs contribution to heritability of traits, a Stratified LD Score Regression (sLDSC) was conducted on UKBB GWAS summary statistics estimated by Neale Lab (http://www.nealelab.is/uk-biobank/) for 50 complex traits. We used 503 samples of European ancestry from the 1000 Genomes Project^53^ to compute LD Scores; HapMap Project Phase 3^54^ SNPs were kept for the analysis, after filtering for minor allele frequency above 1% and INFO score above 95%. 50 TAGR sets were used as functional categories, in addition to another set of 50 “negative TAGRs” defined excluding 500kbs around each GWAS Catalog hit.

### Models tested and covariates

We regressed each trait *t* on a linear model including the standardized *covA* for the ancestry *p* and a vector of covariates **c**:

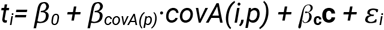

then estimated the *covA* coefficient. Categorical traits, which were transformed to {0, 1} where 1 stands for the category described in **Table S4** and 0 for all the others, were regressed with a logistic regression. We instead adopted an ordinal logistic regression for ordinal traits: the category in **Table S4** here describes the category order.

The covariates included in the model are (in parenthesis the corresponding UKBB Field ID): age (21022), sex (31), age^2^, age×sex, age^2^×sex, latitude (22703), longitude (22701), UKBB assessment centre (54), urban or rural home area (20118, defined collapsing codes 1,5,11,12 into “urban”; 2,6,13,14,15 into “town”; 3,4,7,8,16,17,18 into “rural” categories), qualifications (6138), age when completed education (845), Townsend deprivation index at recruitment (22189), two *covAs* computed genome-wide. Note that given that *covA* is a relative distance, with three ancestries we only have two degrees of freedom so adding the third would have generated perfect multicollinearity. *covA*s were computed on the corresponding TAGRs for each trait, or genome-wide where explicitly mentioned.

Then, the slope coefficient (*β*_*covA*_), or the Odds Ratio (OR_*covA*_) were directly used to assess ancestry-trait association for continuous and categorical traits respectively, and significance from double-sided coefficient p-value from glm R function. Significance was evaluated at Benjamini-Hochberg False Discovery Rate = 0.05. In all cases when reporting results for each trait we count multi-category traits only once.

## Acknowledgements

This research has been conducted using the UK Biobank Resource under Application Number 19655. D.M. was supported by the University of Turin through Grant For Internationalization 2022. V.P. and M.Met. were supported by the European Union through Horizon 2020 Research and Innovation Program under Grant No. 810645 and the European Union through the European Regional Development Fund Project No. MOBEC008. M.Met. was supported by the Estonian Research Council grant PUT (PRG1899). D.F. is a PhD student enrolled in the National PhD in Artificial Intelligence, XXXVIII cycle, course on Health and life sciences, organized by Università Campus Bio-Medico di Roma. We would like to thank Francesco Montinaro and Alessandro Raveane for insightful discussions. Data analyses for this study were carried out in the High-Performance Computing Center of the University of Tartu.

## Author contributions

V.P., L.P. and D.M. conceived and designed the study; V.P. and D.M. developed the computational pipelines and performed the bulk of statistical analyses; M.Mez. contributed to demographic simulations; M.Mez., S.A. and D.F. contributed to data analysis; J.F.W., M.Met., P.P., L.P. provided data and contributed to results interpretation; V.P. and D.M. drafted the manuscript; all authors reviewed and approved the submitted paper.

## Code Availability

Simulations: https://github.com/vasilipankratov/SLiM_quantitative_trait

## References

1. Uffelmann, E. et al. Genome-wide association studies. Nat. Rev. Methods Primer 1, 59 (2021).

2. Abdellaoui, A., Yengo, L., Verweij, K. J. H. & Visscher, P. M. 15 years of GWAS discovery: Realizing the promise. Am. J. Hum. Genet. 110, 179–194 (2023).

3. Bycroft, C. et al. The UK Biobank resource with deep phenotyping and genomic data. Nature 562, 203–209 (2018).

4. Leitsalu, L. et al. Cohort profile: Estonian biobank of the Estonian genome center, university of Tartu. Int. J. Epidemiol. 44, 1137–1147 (2015).

5. Sijtsma, A. et al. Cohort Profile Update: Lifelines, a three-generation cohort study and biobank. Int. J. Epidemiol. 51, e295–e302 (2022).

6. Kurki, M. I. et al. FinnGen provides genetic insights from a well-phenotyped isolated population. Nature 613, 508–518 (2023).

7. Wang, Y. et al. Polygenic prediction across populations is influenced by ancestry, genetic architecture, and methodology. Cell Genomics 100408 (2023) doi:10.1016/j.xgen.2023.100408.

8. Martin, A. R. et al. Clinical use of current polygenic risk scores may exacerbate health disparities. Nat. Genet. 51, 584–591 (2019).

9. Popejoy, A. B. & Fullerton, S. M. Genomics is failing on diversity. Nature 538, 161–164 (2016).

10. Kachuri, L. et al. Principles and methods for transferring polygenic risk scores across global populations. Nat. Rev. Genet. (2023) doi:10.1038/s41576-023-00637-2.

11. Price, A. L. et al. Principal components analysis corrects for stratification in genome-wide association studies. Nat. Genet. 38, 904–909 (2006).

12. Loh, P.-R. et al. Efficient Bayesian mixed-model analysis increases association power in large cohorts. Nat. Genet. 47, 284–290 (2015).

13. Bulik-Sullivan, B. et al. LD score regression distinguishes confounding from polygenicity in genome-wide association studies. Nat. Genet. 47, 291–295 (2015).

14. Zhou, W. et al. Efficiently controlling for case-control imbalance and sample relatedness in large-scale genetic association studies. Nat. Genet. 50, 1335–1341 (2018).

15. Kerminen, S. et al. Fine-Scale Genetic Structure in Finland. G3 GenesGenomesGenetics 7, 3459–3468 (2017).

16. Pankratov, V. et al. Differences in local population history at the finest level: the case of the Estonian population. Eur. J. Hum. Genet. 28, 1580–1591 (2020).

17. O’Dushlaine, C. T. et al. Population structure and genome-wide patterns of variation in Ireland and Britain. Eur. J. Hum. Genet. 18, 1248–1254 (2010).

18. Helgason, A., Yngvadóttir, B., Hrafnkelsson, B., Gulcher, J. & Stefánsson, K. An Icelandic example of the impact of population structure on association studies. Nat. Genet. 37, 90–95 (2005).

19. Zaidi, A. A. & Mathieson, I. Demographic history mediates the effect of stratification on polygenic scores. eLife 9, 1–30 (2020).

20. Sul, J. H., Martin, L. S. & Eskin, E. Population structure in genetic studies: Confounding factors and mixed models. PLOS Genet. 14, e1007309 (2018).

21. Pärna, K. et al. A Principal Component Informed Approach to Address Polygenic Risk Score Transferability Across European Cohorts. Front. Genet. 13, 899523 (2022).

22. Kerminen, S. et al. Geographic Variation and Bias in the Polygenic Scores of Complex Diseases and Traits in Finland. Am. J. Hum. Genet. 104, 1169–1181 (2019).

23. Cook, J. P., Mahajan, A. & Morris, A. P. Fine-scale population structure in the UK Biobank: implications for genome-wide association studies. Hum. Mol. Genet. 29, 2803–2811 (2020).

24. Ding, Y. et al. Polygenic scoring accuracy varies across the genetic ancestry continuum. Nature 618, 774–781 (2023).

25. Mostafavi, H. et al. Variable prediction accuracy of polygenic scores within an ancestry group. eLife 9, e48376 (2020).

26. Barton, N., Hermisson, J. & Nordborg, M. Why structure matters. eLife 8, e45380 (2019).

27. Berg, J. J. et al. Reduced signal for polygenic adaptation of height in UK Biobank. eLife 8, 1–47 (2019).

28. Sohail, M. et al. Polygenic adaptation on height is overestimated due to uncorrected stratification in genome-wide association studies. eLife 8, 1–17 (2019).

29. Lazaridis, I. et al. Ancient human genomes suggest three ancestral populations for present-day Europeans. Nature 513, 409–13 (2014).

30. Haak, W. et al. Massive migration from the steppe was a source for Indo-European languages in Europe. Nature 522, 207–211 (2015).

31. Allentoft, M. E. et al. Population genomics of Bronze Age Eurasia. Nature 522, 167–172 (2015).

32. Marchi, N. et al. The genomic origins of the world’s first farmers. Cell 185, 1842–1859.e18 (2022).

33. Cox, S. L., Ruff, C. B., Maier, R. M. & Mathieson, I. Genetic contributions to variation in human stature in prehistoric Europe. Proc. Natl. Acad. Sci. U. S. A. 116, 21484–21492 (2019).

34. Ju, D. & Mathieson, I. The evolution of skin pigmentation-associated variation in West Eurasia. Proc. Natl. Acad. Sci. 118, e2009227118 (2021).

35. Ali J. Berens, Taylor L. Cooper, & Joseph Lachance. The Genomic Health of Ancient Hominins. Hum. Biol. 89, 7 (2017).

36. Olalde, I. et al. Derived immune and ancestral pigmentation alleles in a 7,000-year-old Mesolithic European. Nature 507, 225–8 (2014).

37. Mathieson, I. et al. Genome-wide patterns of selection in 230 ancient Eurasians. Nature 528, 499–503 (2015).

38. Saag, L. et al. Genetic ancestry changes in Stone to Bronze Age transition in the East European plain. Sci. Adv. 7, eabd6535 (2021).

39. Marnetto, D. et al. Ancestral genomic contributions to complex traits in contemporary Europeans. Curr. Biol. 32, 1412–1419.e3 (2022).

40. Haller, B. C. & Messer, P. W. SLiM 4: Multispecies Eco-Evolutionary Modeling. Am. Nat. 201, E127–E139 (2023).

41. Sakaue, S. et al. Dimensionality reduction reveals fine-scale structure in the Japanese population with consequences for polygenic risk prediction. Nat. Commun. 11, 1569 (2020).

42. Mallick, S. et al. The Allen Ancient DNA Resource (AADR): A curated compendium of ancient human genomes. http://biorxiv.org/lookup/doi/10.1101/2023.04.06.535797 (2023) doi:10.1101/2023.04.06.535797.

43. Sollis, E. et al. The NHGRI-EBI GWAS Catalog: knowledgebase and deposition resource. Nucleic Acids Res. 51, D977–D985 (2023).

44. Finucane, H. K. et al. Partitioning heritability by functional annotation using genome-wide association summary statistics. Nat. Genet. 47, 1228–1235 (2015).

45. Kristal-Boneh, E., Froom, P., Harari, G., Malik, M. & Ribak, J. Summer-Winter Differences in 24 h Variability of Heart Rate. Eur. J. Cardiovasc. Risk 7, 141–146 (2000).

46. De Lorenzo, F. Cold adaptation and the seasonal distribution of acute myocardial infarction. QJM 92, 747–751 (1999).

47. Mourot, L., Bouhaddi, M. & Regnard, J. Effects of the cold pressor test on cardiac autonomic control in normal subjects. Physiol. Res. 83–91 (2009) doi:10.33549/physiolres.931360.

48. LeBlanc, J., Dulac, S., Cote, J. & Girard, B. Autonomic nervous system and adaptation to cold in man. J. Appl. Physiol. 39, 181–186 (1975).

49. Bortolini, E. et al. Early Alpine occupation backdates westward human migration in Late Glacial Europe. Curr. Biol. 31, 2484–2493.e7 (2021).

50. Le, M. K. et al. 1,000 ancient genomes uncover 10,000 years of natural selection in Europe. http://biorxiv.org/lookup/doi/10.1101/2022.08.24.505188 (2022) doi:10.1101/2022.08.24.505188.

51. Baumdicker, F. et al. Efficient ancestry and mutation simulation with msprime 1.0. Genetics 220, iyab229 (2022).

52. Yair, S. & Coop, G. Population differentiation of polygenic score predictions under stabilizing selection. Philos. Trans. R. Soc. B Biol. Sci. 377, 20200416 (2022).

53. The 1000 Genomes Project Consortium et al. A global reference for human genetic variation. Nature 526, 68–74 (2015).

54. The International HapMap 3 Consortium. Integrating common and rare genetic variation in diverse human populations. Nature 467, 52–58 (2010).

