## Supplementary Material for "Ancestral genetic components are consistently associated with the complex trait landscape in European biobanks"

#### Supplementary Methods

##### Simulations

To decrease runtime we split the simulation into three steps described in Table S6.

Each subsequent step takes the output of the previous one as input and inherits the recombination and mutation rates. Where appropriate, multiple instances of subsequent steps started from the same output of the previous step to save run-time and disc space.

Step 1 was performed in msprime v1.2.0[1] using the discrete time Wright-Fisher model[2]. We simulated 1000 unlinked genomic regions each being 20 kb long with a uniform recombination rate of 1e-8 and uniform mutation rate of 1.25e-8 in a panmictic population with a constant effective population size of 14,000. The pyslim v1.0.1 package was used to convert the msprime's tree sequence object into a SLiM sim object.

Step 2 was run in SLiM v4.0[3] simulating a panmictic population of 14,000 diploid individuals. At the very beginning of this step we randomly picked 1000 genetic variants (one per each genomic region) meeting the following criteria: a) maf >= 0.1 at the beginning of step 2; b) position in the genomic region is between 5 and 15 kb. Each of the 1000 variants was then randomly assigned an effect size from either a single or a mixture of two zero-centered normal distributions (Table S7). In the latter case (setup 3 in Table S7) where we simulate lower trait polygenicity (i.e. a small fraction of variants having strong effect on the trait) we sampled the effect sizes from a high variance distribution (a) with probability 0.05 and from a low variance distribution (b) with probability 0.95. Effect sizes are constant across generations and populations hence no gene-by-environment interactions are simulated. All mutations originating during steps 2 and 3 are assumed to have no effect on the trait.

Step 3 was implemented in SLiM v4.0[3]. The modeled demography is described in Figure S10 and Table S8. Specifically, we model the British population as a result of admixture that took place 140 generations ago between Early European Farmers (54%) and Yamnaya (46%). In turn, Early European Farmers are modeled as a result of admixture between Anatolia (78%) and Western Hunter-Gatherers (22%) 270 generations ago. The British population started growing exponentially 17 generations from Ne of 6000 at a rate of 0.18 reaching Ne of approximately 100K by the end of the simulation. Although according to IBD-based estimates[4] the present-day Ne of the British population might be on the order of 10M we aim our simulation at 100K for performance reasons. The sampled ancient genotypes come from reference populations that diverged from the populations actually involved in the admixture events (Anatolia, Western Hunter-Gatherers and Yamnaya) 50 generations before the admixture.

To explore effects of directional selection we modeled trait optima shifts in one of the following populations at a time: Anatolia, Western Hunter-Gatherers, Yamnaya. The optimum was either kept constant at 0 in all populations (no directional selection) or shifted to +0.5 or +1 depending on the specific simulation. The optimum shift is implemented right after the origin of the corresponding population (Table S8). This results in different populations having different amounts of time to adapt to the new optimum.

When population differed in their optima we explored the following approaches to model the trait optimum of admixed populations, resulting in different post-admixture selection regimes:

1. The optimum of an admixed population is a weighted average of the two admixing populations optima. For instance, if the optimum of Anatolia is 1 and the optimum of WHG is 0, the optimum in EFF is 0.78.
2. The optimum of an admixed population is the maximum of the two admixing populations optima. For instance, if the optimum of Anatolia is 1 and the optimum of WHG is 0, the optimum in EFF is 1.
3. The optimum of an admixed population is twice the maximum of the two admixing populations optima. For instance, if the optimum of Anatolia is 1 and the optimum of WHG is 0, the optimum in EFF is 2.
4. The optimum of an admixed population is the minimum of the two admixing populations optima. For instance, if the optimum of Anatolia is 1 and the optimum of WHG is 0, the optimum in EFF is 0.

We additionally modeled trait optimum equal to +2 in the UK or the EFF populations. See Table S9 for more details.

In steps 2 and 3 genetic value for each individual in each generation was calculated by summing up counts of alternative alleles at each causal variant weighted by the corresponding effect size. The trait value is calculated by adding a random variable from a normal zero-centered distribution to the genetic value, mimicking environmental noise (Table S7). The trait is measured in arbitrary units, however, by design its SD is close to 1 (Figure S6).

The fitness of each individual is calculated based on its trait value using the following formula:

$$Fitness = e^{\frac{-(opt-t)^2}{2\times\omega^2}}$$

where *opt* is the trait optimum of the population, *t* is the individual's trait value and $\omega$ the standard deviation of the fitness function; higher $\omega$ values correspond to weaker stabilizing selection.

The latter may take one of the three values: 3, 5 and 10. The choice of those values is inspired by Sanjak et al., 2018[5] who proposed the interquartile range of $\omega$ for present-day


2

human traits to be 5.3 - 13.3 and we assume these values to be lower in ancient populations. Another simulation study[6] used $\omega$ values of 4, 5 and 7 so our range is fully overlapping but wider. All those parameters stay constant across generations (including phase 3) and populations.

As a result of both random assignment of effect sizes and changes in causal variants allele frequencies due to drift, stabilizing and directional selection (in step 3) heritability of the selected trait can differ both between simulations and through time in a given simulation (Table S10).

We sampled 10K genotypes and corresponding trait values from the present-day British population at the end of the simulation and 100 genotypes from each of the three reference populations: Anatolia, WHG and Yamnaya at the time of the corresponding admixture events. The UK samples were pruned to keep only 1 individual out of each pair with first or second degree relatedness, resulting in around 7K individuals remaining.

For CovA calculation we kept only regions with the causal variant effect size meeting the following criteria:

$$\frac{2\beta^2 f(1-f)}{\sigma_t^2} > 1.5\cdot 10^{-5}$$

where $\beta$ is the true effect size on the trait, $f$ is derived allele frequency and $\sigma_t^2$ is the phenotypic variance in the present-day population. The threshold of 1.5e-5 corresponds to an average expected p-value = 5e-8 in a GWAS for a continuous trait with a sample size of 200K. This was done to imitate GWAS discovery and resulted in a median of 495 regions used in covA calculation (min = 351, max = 565).

### Supplementary Figures

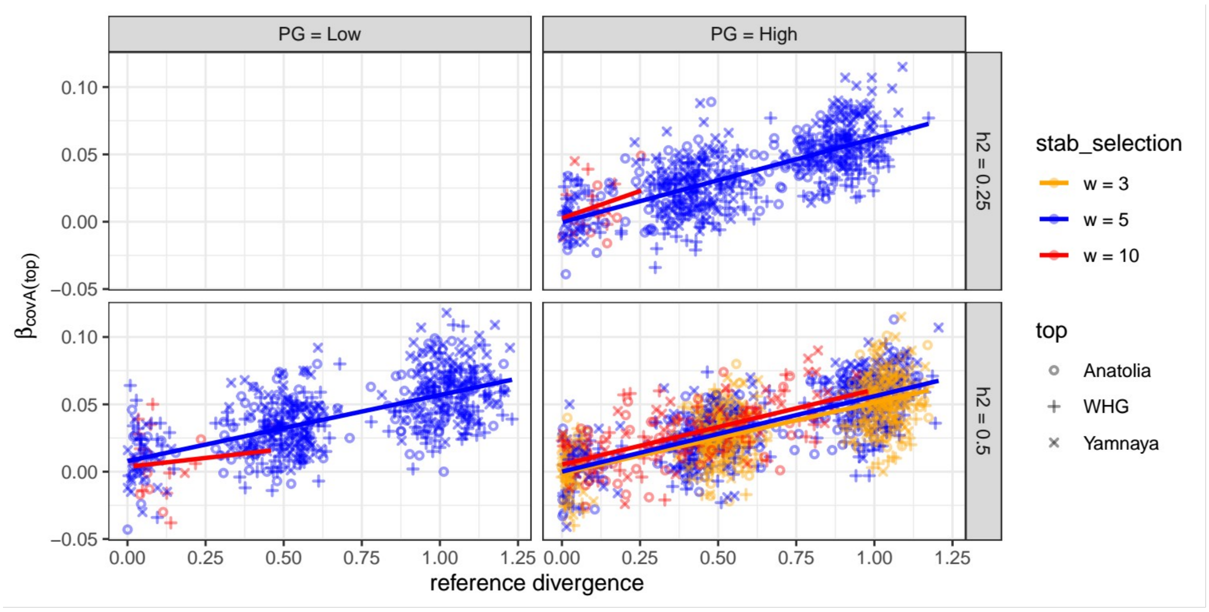


 3

***Supplementary figure 1.*** *Effects of stabilizing selection strength ("stab_selection"), heritability ("h2") and polygenicity ("PG") on the relationship between beta lm(trait~covA) and genetic value differentiation. As in Figure 1 in the main text, each data point corresponds to a single simulation and reports the beta for the ancestry with the highest genetic value ("top"). Scenarios with h2 = 0.25 and PG = low were not tested.*

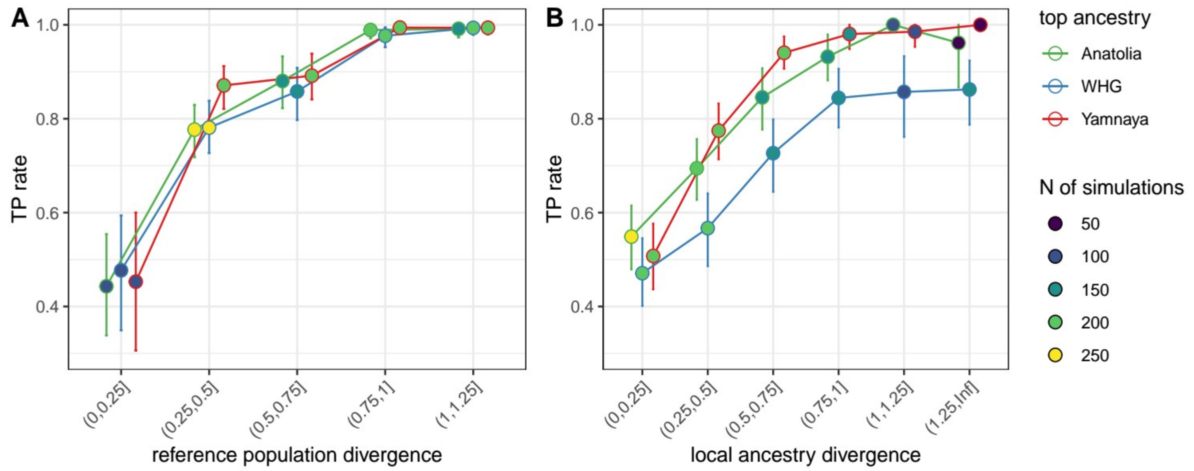


***Supplementary figure 2.*** *Fraction of simulations with the ancestry with the highest $\beta_{covA}$ ('top ancestry') having also the highest mean genetic value among reference populations (A) or local ancestries in the contemporary genomes (B) depending on the level of ancestry differentiation (defined as in Figure 1). The whiskers represent 95% confidence intervals obtained by bootstrapping simulations 1000 times.*

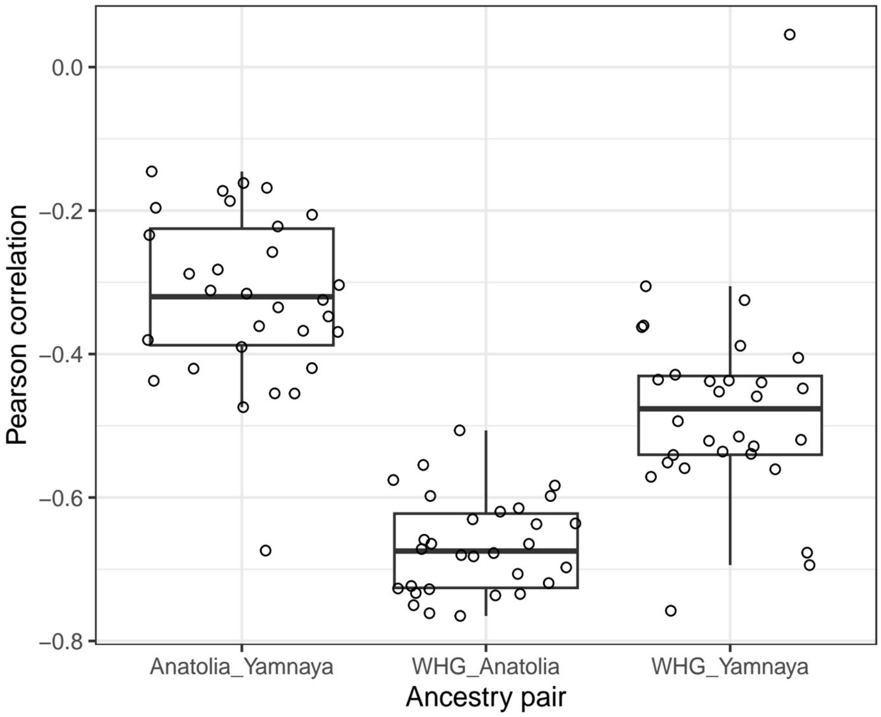


***Supplementary Figure 3.*** *Correlation between $\beta_{covA(j)}$ for different ancestries. Each data point is on the plot is a Pearson's correlation coefficient between $\beta_{covA(j)}$ for the two specified ancestries under a specific simulation scenario. Each correlation estimate is based on at least 80 runs. See table S11 for details.*

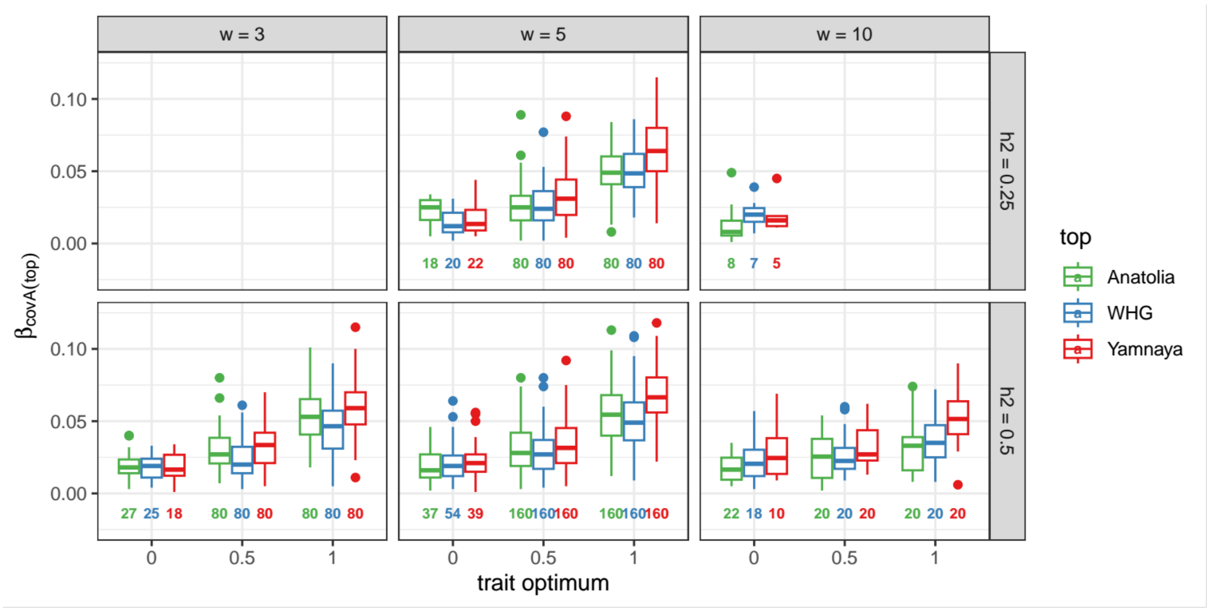


***Supplementary figure 4.*** *Distribution of beta values for the ancestry with the highest genetic value (which always matches the ancestry that underwent a shift in the trait optimum in scenarios where such shift was modeled) in scenarios with all three reference populations having the trait optimum at zero ("0" on x-axis) or one of the ancestries experiencing an increase of the optimum by 0.5 or 1 arbitrary units. For reference, trait sd in the contemporary*

*population is mostly within the range between 0.9 and 1 (Figure S0). The boxes show $25^{th}$, $50^{th}$ and $75^{th}$ quantiles, while the whiskers show values within 1.5 times the interquartile range (IQR).*

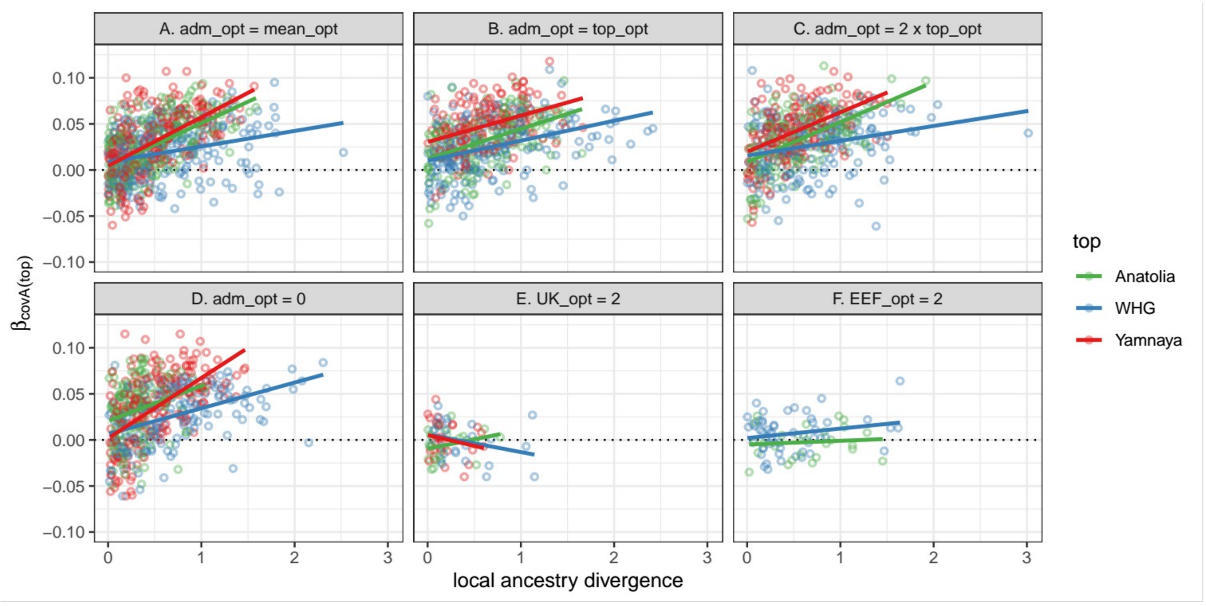


***Supplementary figure 5.*** *Relationship between lm(trait~covA) beta and local ancestry divergence, defined as the difference between the highest and the second highest mean genetic values among the three local ancestries, normalized by trait standard deviation in the present-day population. See Supplementary Methods for a detailed description of the different scenarios of post-admixture selection (A-F).*

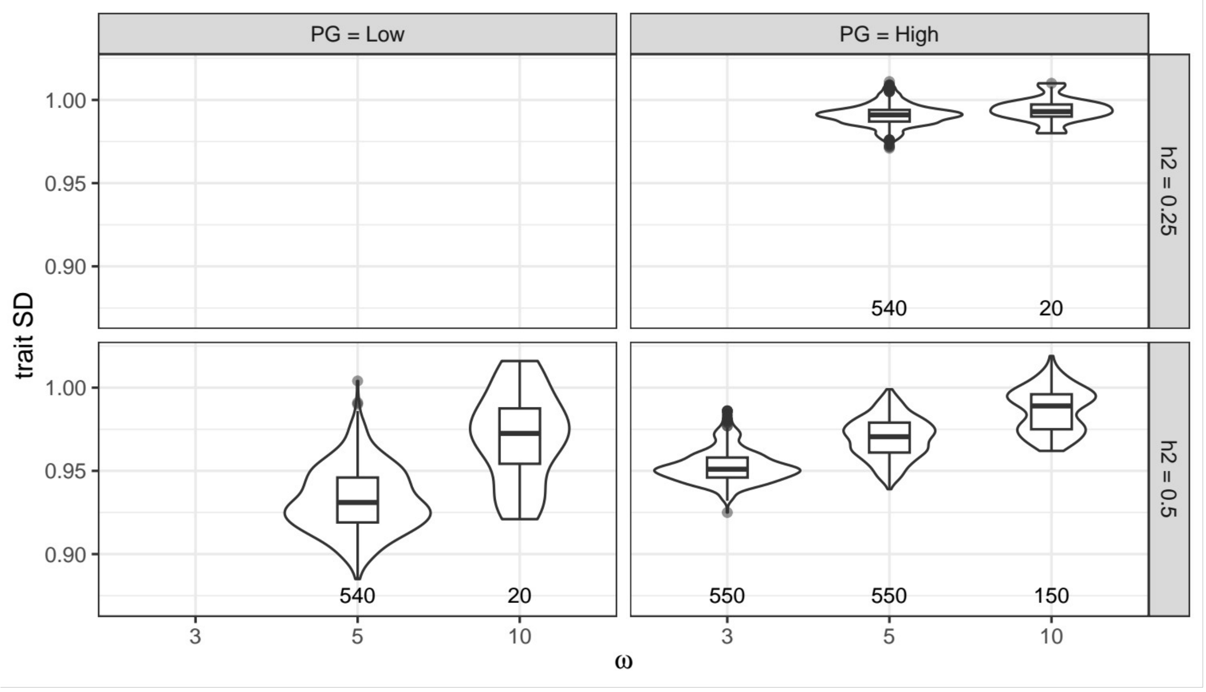



6

**Supplementary Figure 6.** *Trait standard deviation in the contemporary population for various values of polygenicity (PG), heritability (h2) and stabilizing selection (expressed as the standard deviation of the fitness function, w). Numbers below the boxplots show the number of simulations in each category. Scenarios with h2=0.25 and PG=Low were not tested. The boxes show 25th, 50th and 75th quantiles, while the whiskers show values within 1.5 times the interquartile range (IQR).*

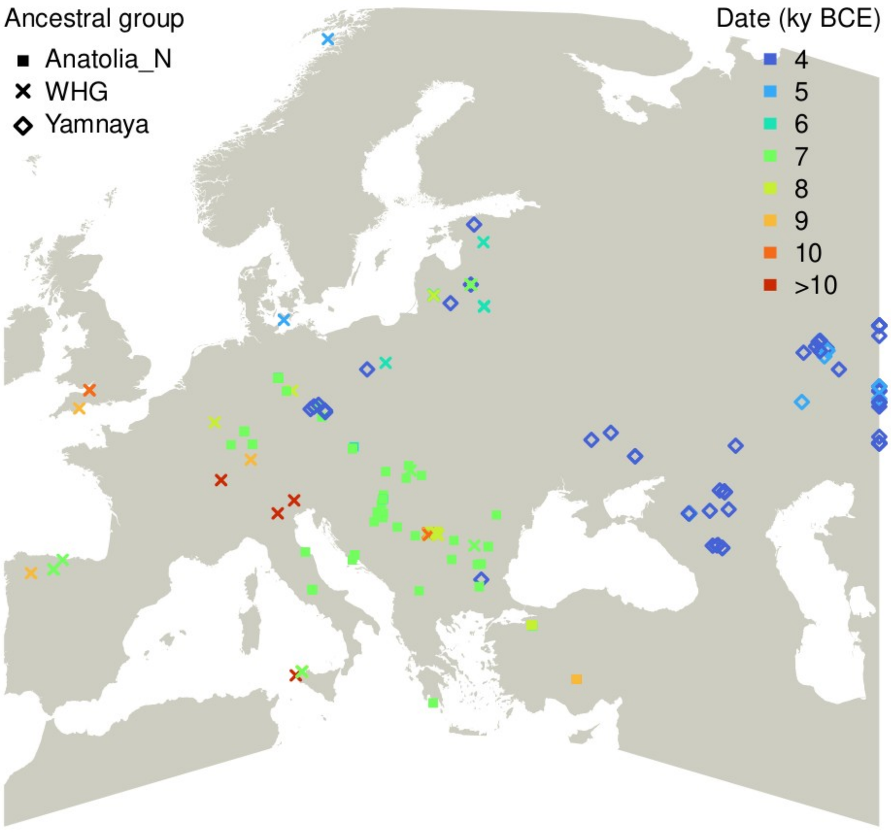


**Supplementary Figure 7.** *Geographical locations of samples assigned to the three ancestral groups considered, with color coding for their average date estimate in thousand years Before Current Era.*

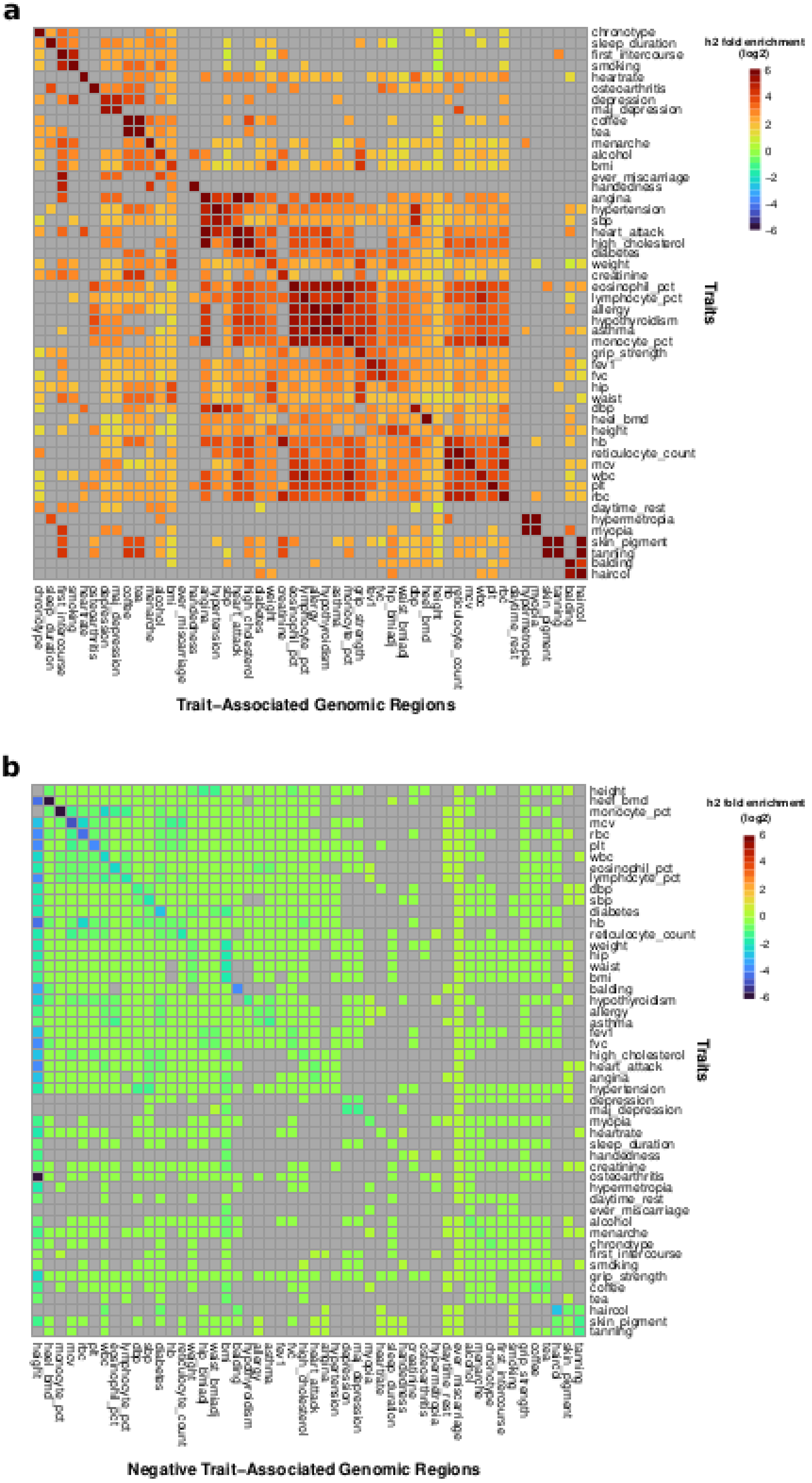


***Supplementary figure 8.*** *Heritability enrichment estimates for (A) TAGRs (50 annotations), and (B) negative TAGRs, defined excluding GWAS hits.. Enrichment of traits expressed as (Proportion of heritability) /(Proportion of SNPs). All colored cells indicate nominal significance at P<0.05, while gray cells are not significant. Annotations are ordered by clustering. Overall, all traits show their highest enrichment in correspondent genomic regions.*


8

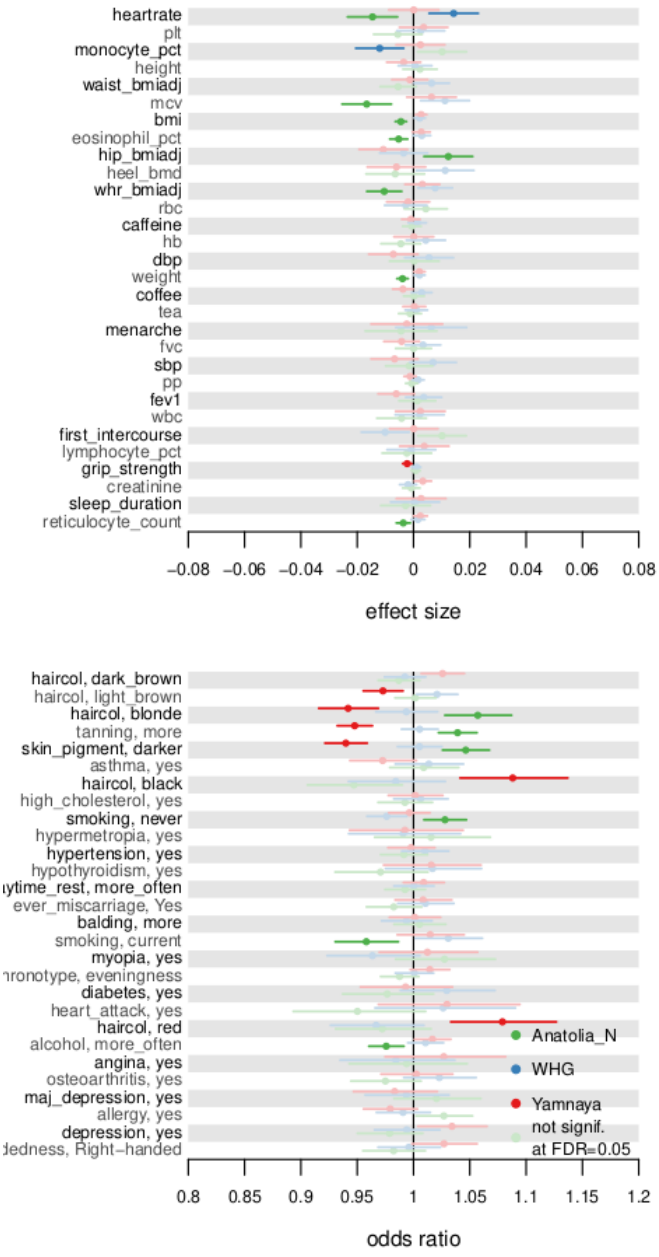


***Supplementary Figure 9.*** *Trait-ancestry associations in UKBB using genome-wide covAs. Besides the origin of the covA metric, the graph details correspond to what shown in Figure 2A.*

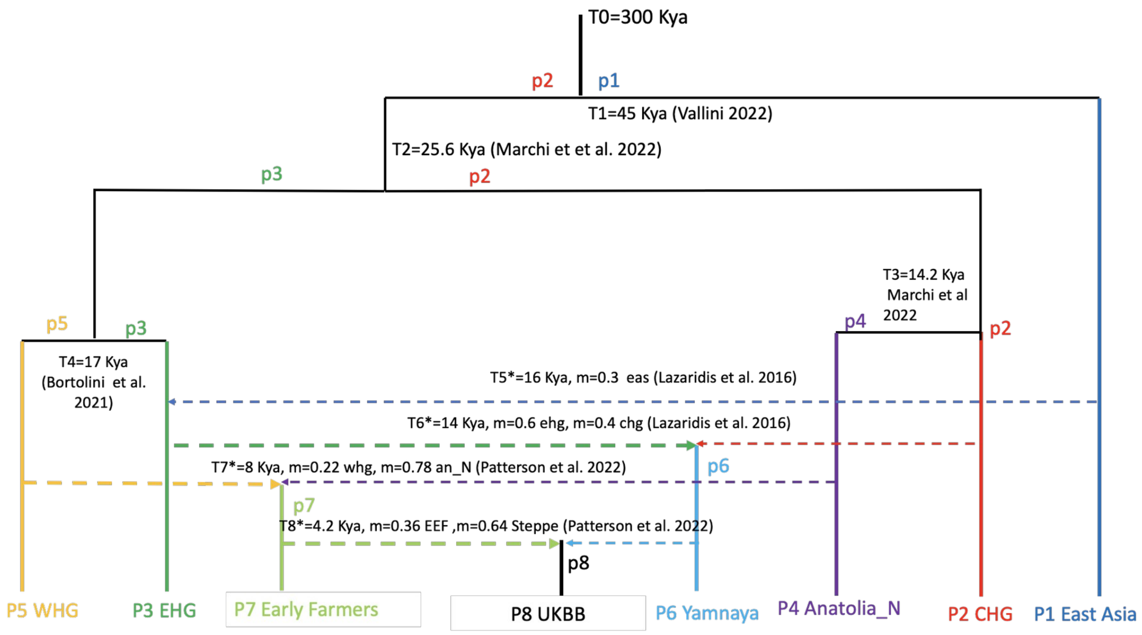


***Supplementary Figure 10.*** *In this graph we report the demographic scenario used for the simulation framework. Admixture and migration events are characterized by the symbol * . The code for the populations is the following: WHG (Western Hunther gathers), EHG (Eastern Hunter gathers), EEF or Early farmers, UKBB (UK Biobank present-day individuals), Yamnaya or Steppe, Anatolia_N (Anatolia Neolithic), CHG (Caucasus Hunter gathers) and East Asia. In the graph we report the estimated time for each event (split, admixture, migration) measured in thousands of years according to the literature, as follows: T1 - Vallini et al 2022[7]; T2,T3 Marchi et al 2022[8]; T4 - Bortolini et al 2021[9] ; T5,T6 - Lazaridis et al 2016[10]; T7,T8 - Patterson et al. 2022[11]*


10

### Supplementary Tables

***Supplementary Table 1.*** *$\beta_{covA}$ confidence intervals in scenarios with all three ancestries having the same trait optima. To explore the variability in the results due to pulling together runs with different ω and h2 obtained the CI for different simulation groups. Each simulation group includes all simulation runs with specific combinations of ω and h2 listed in the second column.*

| *Simulations group* | *Combined scenarios (based on ω and h2)* | *N of simulations* | *q2.5* | *q97.5* |
|---|---|---|---|---|
| *1* | *w=3; h2=0.5*<br>*w=5; h2=0.25*<br>*w=5; h2=0.5*<br>*w=10; h2=0.25*<br>*w=10; h2=0.5* | *330* | *0.0030* | *0.0483* |
| *2* | *w=5; h2=0.25*<br>*w=5; h2=0.5*<br>*w=10; h2=0.25*<br>*w=10; h2=0.5* | *260* | *0.0030* | *0.0500* |
| *3* | *w=5; h2=0.25*<br>*w=5; h2=0.5*<br>*w=10; h2=0.5* | *240* | *0.0030* | *0.0501* |
| *4* | *w=5; h2=0.25*<br>*w=5; h2=0.5* | *190* | *0.0030* | *0.0471* |
| *5* | *w=10; h2=0.25*<br>*w=10; h2=0.5* | *70* | *0.0037* | *0.0519* |

***Supplementary Table 2.*** *Simulation results (see supplementary spreadsheet).*

***Supplementary Table 3.*** *Classification of ancient samples into ancestral group, together with geographical coordinates, dates and PCs used for classification (see supplementary spreadsheet).*

***Supplementary Table 4.*** *Traits description, sample sizes and transformation (see supplementary spreadsheet).*

***Supplementary Table 5.*** *GWAS Catalog search terms for the definition of TAGRs (see supplementary spreadsheet).*

***Supplementary Table 6.*** *Simulation steps.*


11

| Step | Simulator | Duration | Description |
|---|---|---|---|
| 1. Neutral burn-in | msprime | Till coalescence | Generating genetic diversity at mutation-drift equilibrium |
| 2. Burn-in with stabilizing selection | SLiM | 1000 generations | Picking causal variants, assigning effect sizes to those and equilibrating genetic variance to stabilizing selection |
| 3. Main simulation | SLiM | 1500 generations | Simulating all demographic events, directional selection and sampling genomes and phenotypes |

***Supplementary Table 7.*** *Distributions used to draw variant effect sizes and environmental noise values.*

| Setup | Polygenicity | Target h2 | Effect size variance | Environmental noise variance |
|---|---|---|---|---|
| 1 | High | 0.5 | 1.5/1000 | 0.5 |
| 2 | High | 0.25 | 0.75/1000 | 0.75 |
| 3 | Low | 0.5 | a) 0.5*1.5/50 with prob 0.05<br>b) 0.5*1.5/950 with prob 0.95 | 0.5 |

***Supplementary Table 8.*** *Description of the simulated demographic history during step 3 of the simulation (see supplementary spreadsheet).*

***Supplementary Table 9.*** *Number of simulation runs for each parameter set (see supplementary spreadsheet).*

***Supplementary Table 10.*** *Trait heritability observed in the simulations.*

| Target h2 | $\omega$ | Polygenicity | Median h2 (min-max) at the end of step 2 | Median h2 (min-max) at the end of step 3 |
|---|---|---|---|---|
| 0.25 | 5 | High | 0.256 (0.234-0.276) | 0.237 (0.204-0.266) |
| 0.25 | 10 | High | 0.259 (0.245-0.280) | 0.242 (0.223-0.268) |
| 0.5 | 3 | High | 0.490 (0.461-0.519) | 0.448 (0.419-0.488) |
| 0.5 | 5 | High | 0.495 (0.470-0.520) | 0.469 (0.414-0.503) |

 12

| | | | | |
|---|---|---|---|---|
| 0.5 | 5 | Low | 0.478 (0.400-0.559) | 0.422 (0.367-0.506) |
| 0.5 | 10 | High | 0.508 (0.485-0.548) | 0.488 (0.459-0.516) |
| 0.5 | 10 | Low | 0.507 (0.458-0.552) | 0.471 (0.412-0.52) |

### References


1. Baumdicker, F. *et al.* Efficient ancestry and mutation simulation with msprime 1.0. *Genetics* **220**, iyab229 (2022).
2. Nelson, D. *et al.* Accounting for long-range correlations in genome-wide simulations of large cohorts. *PLOS Genet.* **16**, e1008619 (2020).
3. Haller, B. C. & Messer, P. W. SLiM 4: Multispecies Eco-Evolutionary Modeling. *Am. Nat.* **201**, E127–E139 (2023).
4. Gilbert, E. *et al.* The Newfoundland and Labrador mosaic founder population descends from an Irish and British diaspora from 300 years ago. *Commun. Biol.* **6**, 469 (2023).
5. Sanjak, J. S., Sidorenko, J., Robinson, M. R., Thornton, K. R. & Visscher, P. M. Evidence of directional and stabilizing selection in contemporary humans. *Proc. Natl. Acad. Sci.* **115**, 151–156 (2018).
6. Yair, S. & Coop, G. Population differentiation of polygenic score predictions under stabilizing selection. *Philos. Trans. R. Soc. B Biol. Sci.* **377**, 20200416 (2022).
7. Vallini, L. *et al.* Genetics and Material Culture Support Repeated Expansions into Paleolithic Eurasia from a Population Hub Out of Africa. *Genome Biol. Evol.* **14**, evac045 (2022).
8. Marchi, N. *et al.* The genomic origins of the world's first farmers. *Cell* **185**, 1842-1859.e18 (2022).
9. Bortolini, E. *et al.* Early Alpine occupation backdates westward human migration in Late Glacial Europe. *Curr. Biol.* **31**, 2484-2493.e7 (2021).
10. Lazaridis, I. *et al.* Genomic insights into the origin of farming in the ancient Near East. *Nature* **536**, 419–424 (2016).
11. Patterson, N. *et al.* Large-scale migration into Britain during the Middle to Late Bronze Age. *Nature* **601**, 588–594 (2022).


 13
